# Transcriptome analysis reveals infection strategies employed by *Fusarium graminearum* as a root pathogen

**DOI:** 10.1101/2020.05.17.092288

**Authors:** Yi Ding, Donald M Gardiner, Kemal Kazan

**Affiliations:** Agriculture and Food, Commonwealth Scientific and Industrial Research Organization, 306 Carmody Road, St Lucia, 4067, Queensland, Australia; Queensland Alliance for Agriculture and Food Innovation (QAAFI), Alliance for Agriculture and Food Innovation (QAAFI), The University of Queensland, Brisbane, St Lucia, 4067, Queensland, Australia

## Abstract

The fungal pathogen *Fusarium graminearum* infect both heads and roots of cereal crops causing several economically important diseases such as head blight, seedling blight, crown rot and root rot. Trichothecene mycotoxins such as deoxynivalenol (DON), a well-known virulence factor, produced by *F. graminearum* (*Fg*) during disease development is also an important health concern. Although how *F. graminearum* infects above-ground tissues is relatively well studied, very little is known about molecular processes employed by the pathogen during below-ground infection. Also unknown is the role of DON during root infection. In the present study, we analyzed the transcriptome of *F. graminearum* during root infection of the model cereal *Brachypodium distachyon.* We also compared our *Fg* transcriptome data during root infection with those reported during wheat head infection. These analyses suggested that both shared and unique infection strategies employed by the pathogen during colonization of different host tissues. Several metabolite biosynthesis genes induced in *F. graminearum* during root infection could be linked to phytohormone production, implying that the pathogen likely interferes root specific defenses. In addition, to understand the role of DON in *Fg* root infection, we analyzed the transcriptome of the DON deficient *Tri5* mutant. These analyses showed that the absence of DON had a significant effect on fungal transcriptional responses. Although DON was produced in infected roots, this mycotoxin did not act as a virulence factor during root infection. Our results reveal new mechanistic insights into the below-ground strategies employed by *F. graminearum* that may benefit the development of new genetic tools to combat this important cereal pathogen.

## Introduction

Fungal plant pathogens have adopted versatile strategies to colonize their hosts. While some fungal pathogens show strict host and tissue specificity, others can adjust their lifestyles to infect different hosts and tissues. Some fungal pathogens such as the rice blast *Magnaporthe oryzae* (Marcel et al., 2010; Sesma and Osbourn, 2004) and the corn smut *Ustilago maydis* (Mazaheri-Naeini et al., 2015), which commonly invade above-ground plant parts, can also undergo developmental processes resembling to root infecting fungi. Such changes in the pathogen may require sensing of host signals and we previously showed that sensing of root signals prior to root infection by *Fg* can indeed lead to developmental changes in the pathogen (Ding et al., 2020). As a member of the *Fusarium* species complex, *Fusarium graminearum* (*Fg*) causes Fusarium Head Blight (FHB) or scab, one of the most economically important diseases of cereal crops. FHB causes substantial yield losses and mycotoxin contaminations of grains, resulting in billions of dollars of economic losses worldwide and threatening our food supply and safety (Chen et al., 2019; Trail, 2009). Most studies on FHB have so far focused on wheat heads as the pathogen initially infects individual wheat florets from which it can spread to other florets through the rachis and can eventually colonize the whole spike. However, recent research has shown that *Fg* is also capable of infecting roots and young seedlings of wheat, barley and maize, causing crown rot, root rot and seedling blight (Henkes et al., 2011; Lanoue et al., 2010; Stephens et al., 2008; Wang et al., 2015; Zhou et al., 2019). During the initiation of root infection, *Fg* forms a peg structure outside the root surface and move inter- and intra-cellularly without causing root necrosis. This early colonization stage is followed by a transition of the fungus to a necrotrophic life style where lesions develop and spread to stems and aboveground tissues (Wang et al., 2015).

The availability of the complete genome sequence of *Fg* makes investigations of global regulation of gene expression in this fungus feasible (Kazan and Gardiner, 2018a; Ma et al., 2013). Infection strategies of *Fg* evaluated by transcriptome analyses in different hosts and tissues exclusively during infection of their above-ground tissues such as heads, stems and coleoptiles revealed mostly distinct, but also common gene expression patterns (Boedi et al., 2016; Harris et al., 2016; Lysøe et al., 2011; Zhang et al., 2012, 2016). Interestingly, different cereal species can produce different defense-related metabolites (Dutartre et al., 2012) and such differences may explain why *Fg* might need to tailor its arsenal during colonization of different hosts (Harris et al., 2016). A recent comparative transcriptomic study of *Fg* also showed differential expression of fungal genes during infection of FHB resistant and susceptible wheat genotypes (Pan et al., 2018).

Although transcriptome studies have provided clues associated with host specificity of the *Fg* infection process, fungal transcriptomes of *Fg* mutants with altered virulence have rarely been tested on the same hosts. The mycotoxin deoxynivalenol (DON) is a *Fg* virulence factor during infection of wheat heads (Proctor et al., 1995). DON may also be needed during the interaction of *Fg* with its broader environment (Audenaert et al., 2013). The first step in DON biosynthesis is catalyzed by the trichodiene synthase Tri5, which cyclizes farnesyl pyrophosphate to trichodiene (Hohn and Beremand, 1989). In contrast to its strong expression pattern during wheat head infection, *Tri5* did not show increased *in planta* expression during wheat coleoptile infection, suggesting that DON’s effect on pathogen virulence is tissue specific (Zhang et al., 2012). Other studies have suggested a crucial role for DON during the colonization of wheat stems by *Fg* and the related pathogens *F*. *pseudograminearum* and *F. culmorum* (Desmond et al., 2008; Mudge et al., 2006; Powell et al., 2017; Scherm et al., 2013a). The recent finding where Fhb7-mediated FHB and crown rot disease resistance relies on DON detoxification also highlighted virulence function of this mycotoxin in wheat (Wang et al., 2020). Furthermore, DON is known to activate defense gene expression in wheat (Desmond et al., 2008). Indeed, host transcriptional changes observed in *Brachypodium distachyon* (*Bd*) and wheat spikelets infected by the *Tri5* deletion mutants (ΔTri5) differed from those by wildtype *Fg* (Brauer et al., 2020; Pasquet et al., 2014). DON biosynthesis in *Fg* is regulated by *Tri6* and *Tri10* transcription factors. Analyses of deletion mutants for these genes by transcriptome profiling during plant infection revealed significant transcriptional alterations for a large number of genes, many of which have not been implicated previously in toxin production (Seong et al., 2009). Genetic analyses undertaken in *Fg* have identified many genes influencing DON biosynthesis (Chen et al., 2019). However, DON-non-producing mutants have not been employed for evaluating the effect of this toxin on global transcriptional responses in *Fg*.

Phytohormones mediate immune responses in plants after pest or pathogen attack (Pieterse et al., 2009). In turn, plant pathogenic fungi have evolved ways to compromise host hormone pathways. This is achieved by degrading or producing phytohormones or interfering with their signaling pathways (Kazan and Lyons, 2014; Patkar and Naqvi, 2017). For instance, emerging evidence suggests that phytohormones such as abscisic acid (ABA), gibberellic acid (GA) and ethylene (ET) produced by fungi participate in pathogenicity (Chanclud and Morel, 2016). Previous studies indicated that *Fg* can likely produce auxin (IAA) and ET that may be utilized for attenuating host defenses during FHB (Foroud et al., 2019; Luo et al., 2016; Svoboda et al., 2019). In addition, SA hydroxylases were proposed to be involved in the degradation of host SA by *Fg* (Hao et al., 2019; Qi et al., 2019; Rocheleau et al., 2019). However, how hormonal compounds produced by *Fg* or the host plant are metabolized or involved in host infection is poorly studied. This is at least in part due to potential co-existence of phytohormones derived from both host and the pathogen in the infected tissue and the lack of knowledge on fungal genes involved in phytohormone biosynthesis.

Currently, potential molecular mechanisms employed by *Fg* during root infection are unknown (Kazan and Gardiner, 2018a). A global transcriptome analysis would provide a powerful way to broadly reveal previously unknown features during root infection, thus promoting development of new strategies for combating this pathogen. In addition, to what extent DON may affect global transcription in the pathogen has not been investigated. We previously reported an RNA-seq based transcriptome profiling of *Fg* prior to its physical contact with *Bd* roots. This analysis enabled us to discover novel genes that are involved in nitric oxide (NO) production in *Fg* upon sensing of root signals (e.g. metabolites found in the root exudates) and pathogen virulence (Ding et al., 2020). In this study, using *Bd* as a cereal model, we asked how *Fg* behaves as a root pathogen. To answer this question, we analyzed the transcriptome of *Fg* during infection of *Bd* roots. We analyzed phytohormone levels of infected *Bd* roots to understand potential roles played by phytohormones derived from the fungus. In addition, by comparing the transcriptomes of WT *Fg* and the DON deficient *Tri5* mutant, we uncovered novel insights into global effects of DON on fungal gene expression and metabolism during the infection of host roots.

## Materials and methods

### Plant and fungal materials and root infection assay

The *Fg* CS3005 WT, *Tri5* mutant (Desmond et al., 2008) and *Tri5-GFP* expressing (Gardiner et al., 2009) strains were routinely maintained on Potato Dextrose Agar (PDA, BD Difco). *Bd* (Bd21-3) seeds were surface sterilized and pre-germinated on filter paper (Whatman) placed in 150mm x 25 mm petri dishes (Corning) for 5 days. *Bd* roots inoculation with the *Fg* strains was carried out as described previously (Ding et al., 2020). Briefly, agar plugs (0.25 cm diameter) taken from fungal culture plates (Carboxymethylcellulose agar) were transferred to center of minimum media (pH 7) plates and pre-grown for 3 days. Five-day-old *Bd* seedlings were placed above the fungal colonies and inoculated for additional 5 days. Three biological replicates for *Fg* WT and ΔTri5 inoculated seedlings, and the *Fg* WT alone mycelia were produced by pooling materials from 10-12 plants or fungal mycelia.

### RNAseq and transcriptomic analyses

Fungal and root materials were frozen in liquid nitrogen immediately after harvest. Total RNA was extracted from homogenized samples using a Qiagen RNeasy plant RNA extraction kit with on-column DNase I (Qiagen) digestion following manufacturer’s instructions. RNA was quantified and quality-checked prior to sequencing. An Illumina HiSeq2500 High Output platform was used to generate 50-base pair single-end reads (Australian Genome Research Facility). Reads quality control, alignment, transcript abundance and differential expression (DE) analyses were performed according to the method described previously (Ding et al., 2020). For pairwise comparison (*Fg*WT-only vs. *Fg*WT-*Bd*, or *Fg*WT-*Bd* vs. ΔTri5-*Bd*), different sample files were normalized and merged. Reads were measured as FPKM (Fragments Per Kilobase of gene model per Million reads mapped), and a normalization method developed by Hart et al. (2013) was used and to eliminate background noise of FPKM values where genes with a Gaussian-fit derived log_2_(FPKM) value higher than -3 were considered as expressed. |log2 fold change| ≥ 1 and Benjamini and Hochberg-adjusted P value < 0.05 were applied to DE genes. RNAseq data are available at NCBI under the accession no. PRJNA631873.

### Annotation and functional categorization of differentially expressed genes (DEGs)

BLAST2GO (Götz et al., 2008) was used to assign annotations for fungal DEGs. BLASTP reciprocal best hit analyses were performed in order to identify putative orthologous genes and match unique gene identifiers of the *Fg* CS3005 and PH-1 strains (Gardiner et al., 2014). Based on the PH-1 identifiers, classification ontology of DEGs from pairwise comparisons was annotated with FungiFun2 and subsequently subjected to enrichment analyses (https://elbe.hki-jena.de/fungifun). Functional categories were considered as enriched in the genome if an enrichment Benjamini-Hochberg adjusted P value is smaller than 0.05. Prediction of protein cellular localizations, secretome and putative effectors, transporters, carbohydrate-active enzymes, lipases, secondary metabolism enzymes and transcription factors was according to previously described methods (Ding et al., 2020).

### Identification of homologous genes between *Fg* and *Fp*

All RNAseq reads were mapped to the *Bd* reference genome first. Unmapped reads were extracted and aligned to the *Fg* CS3005 and *Fp* CS3096 reference genomes, respectively, as per previously described (Ding et al., 2020). Only reads that could be mapped to both genomes were retained. Next, read counts measured as FPKM values were log transformed, normalized and subjected to PCA analysis (Fig. S4). Homologous genes within *Fg* and *Fp* were identified using a reciprocal best BLAST hit (RBBH) approach. Gene orthologs with identity of equal or higher than 99% were kept and used for syntenic analysis using a R package shinyCircos (Yu et al., 2018).

### cDNA synthesis and quantitative real-time PCR analysis

0.5-1 µg total RNA was prepared for first-strand cDNA synthesis using the superscript IV synthesis kit (Invitrogen, USA) and quantitative real-time RT-PCR (qRT-PCR) was performed using the ViiA 7 real-time PCR detection platform (Applied BioSystems). Primers were based on previous studies (Ding et al., 2020; Voigt et al., 2005). Expression levels were normalized to the fungal house-keeping gene α*-Tubulin* and were averaged over three biological replicates.

### Root sectioning and fluorescence microscopy

Root dissection was carried out following a previous described method (Ursache et al., 2018). Briefly, roots were harvested at 5dpi and fixed with 4% paraformaldehyde (Sigma) in PBS buffer (pH=6.9) overnight, washed twice with PBS buffer and cleared with ClearSee solution (Ursache et al., 2018) for 3 days. After clearing, roots were hand sectioned from 3 cm above the tips and stained with 0.2% Basic Fuchsin (Sigma). To image GFP and Basic Fuchsin fluorescence, root samples were observed using 488-nm excitation / 519-nm emission, and 561-nm excitation / 625-nm emission, respectively, on a Zeiss Axio Imager M2 microscopy.

### UHPLC quantification of metabolites

For metabolite extractions, mock and *Fg*-infected roots, fungal mycelia and media samples were collected at 5 dpi, immediately frozen and ground in liquid nitrogen. 100 mg of fine-ground materials were resuspended in 2 mL extraction buffer (Ethyl acetate: methanol: dichloromethane, 3:2:1 v/v). 5 μL extracts from 6 replicates of each conditioned sample were injected into a Waters ACQUITY ARC UPLC system with Photodiode Array and passed through a Phenomenex Kinetex column (C18, 1.7 μm, 100 × 2.1 mm). The mobile phases consisted of solvent A (10 mM ammonium formate in water) and solvent B (10 mM ammonium formate in acetonitrile). The gradient program was a linear gradient from 2-40% solvent B delivered over 22 min followed by 40-80% B over 1.5 min at a constant flow rate of 0.4 mL per minute. As external standards, jasmonic acid (Sigma, 10 mM), methyl-jasmonate (Sigma, 40 mM), salicylic acid (Sigma, 10 mM), deoxynivalenol (Sigma, 100 ng/μL), gibberellic acid (Sigma, 10 mM), indole-3-actic acid (Sigma, 2 mM) were freshly prepared for serial dilutions (10x, 100x, 1000x, 10000x). Metabolites were quantified according to the concentration-gradient derived standard curves of external standards. Chromatography absorbance data were aligned using the following wavelengths: 204nm (for SA, JA and MeJA), 254nm (for IAA and DON) and 214nm (for GA), and extracted using the Empower 3 software (Waters).

## Results and discussion

### The *Fg* transcriptome during *Bd* root infection

Despite various studies investigating the transcriptome of *Fg* during colonization of above-ground tissues, *Fg* transcriptome during root infection has not been studied before (Kazan and Gardiner, 2018a). Therefore, here, we first investigated the *Fg* transcriptome during root infection of the model host *Bd*. This transcriptome experiment, which was initially designed for the discovery of a novel host-sensing mechanism in *Fg* prior its physical contact with roots, included three replicates of each of 1) *Fg* grown on minimal media (MM) (*Fg-*only) 2) *Fg*WT and *Bd* roots grown together on MM without physical contact, 3) WT *Fg* colonizing *Bd* roots on MM (*Fg*WT colonization) and 4) a *Tri5* mutant colonizing *Bd* roots (ΔTri5 colonization). Previously, by comparing 1 with 2, we discovered new regulators involved in host sensing-mediated NO production (Ding et al., 2020). Here, we report on the comparisons between 1 and 3, 1 and 4 and 3 and 4, as detailed analyses of these have not been reported previously.

A total of 174,399,800 single-end reads, including 114,577,185 reads from *Fg* alone and *Fg-*infected *Bd* root samples (Ding et al., 2020) and 59,822,615 reads from the ΔTri5 root samples were generated by Illumina sequencing of mRNA libraries (Suppl. data 1). Prior to read mapping to the *Fg* CS3005 genome (Gardiner et al., 2014), reads aligned to the *Bd* genome were filtered. Of these filtered reads across all the samples, at least 91%, ranging from 5 to 19.9 million, were mapped to the *Fg* reference genome. Among these mapped reads, less than 0.3% matched to multiple genomic locations (Suppl. data 1). Out of the 12590 transcripts detected, 11942, 11291 and 11327 were found actively expressed in *Fg-* only, *in* WT *Fg* colonization and in *Tri5* mutant colonization conditions, respectively.

Our analysis revealed a total of 2049 genes that were differentially expressed (DE) (log2 FC ≥ 1 and adjusted p value < 0.05) (Fig. 1A, Suppl. data 2) during colonization of *Bd* roots relative to *Fg-*only. Of these, 1281 and 768 were up- and down-regulated, respectively, during root colonization. The proportion of DE genes (DEGs) (around 20% of total number of genes found in the *Fg* genome) was similar to those observed during above-ground infection by *Fg* in other studies (Brown et al., 2017; Puri et al., 2016). A previous study indicated that some defense genes are similarly regulated in wheat roots and spikes in response to *Fg* (Q. Wang et al., 2018), suggesting that common strategies might be employed by *Fg* to infect different tissue types. Indeed, we previously reported that fungal knockouts of several DEGs identified here showed defects in both root and head infection (Ding et al., 2020).

**Figure 1.**
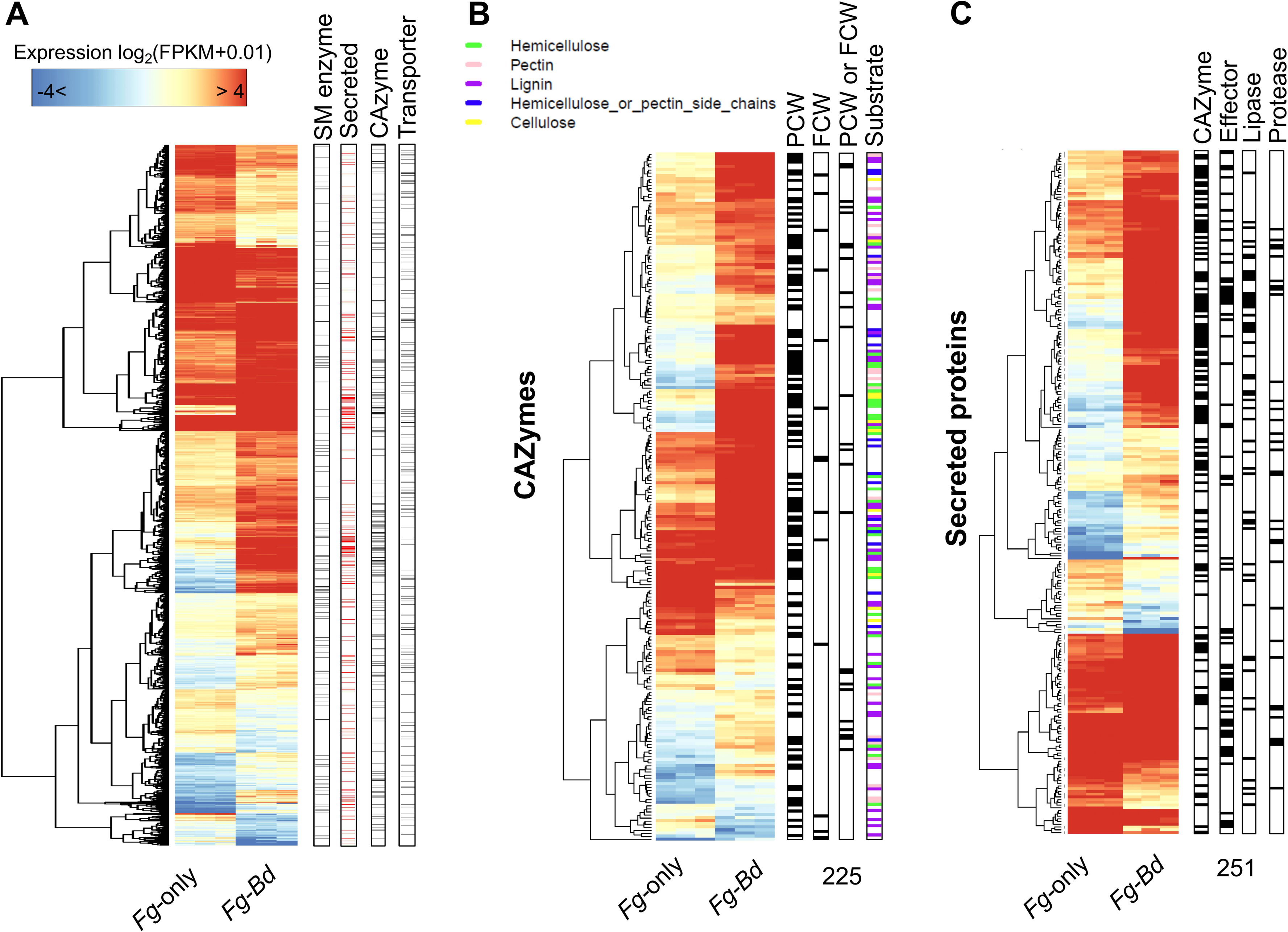
Global regulation of *Fg* genes during *Bd* root colonization. (A) Expression of 2049 *Fg* genes that were differentially regulated during *Bd* root infections (*Fg-Bd*) relative to *Fg* grown in culture (*Fg*-only). Proteins encoded by differentially expressed genes (DEGs) are assigned to four functional categories as shown in the right side of the figure. (B) Key features of 225 DEGs associated with carbohydrate active enzymes (CAZymes). Enzymatic functions and substrate specificities were predicted as shown in the right side of the figure. PCW: plant cell wall, FCW: fungal cell wall. (C) Key features of 251 DEGs encoding putative secreted proteins. *Bd* roots colonized by *Fg* (*Fg-Bd*) were harvested at 5 dpi and *Fg* mycelia grown without *Bd* roots were used as control (*Fg*-only). Transcript levels of fungal genes were presented as normalized FPKM (Fragments Per Kilobase of transcript per Million mapped reads) values and log2-transformed. Heatmap colour range represents high to low expression levels. The dendrogram shows distance similarity of expression of each gene.

### Fungal processes employed by *Fg* during root infection

Detailed analysis of DEGs (*Fg-*only *vs* WT *Fg* colonization) revealed a number of enriched functional categories likely to be used by the pathogen during root infection. Below, some of these functional categories were discussed in more detail.

### Genes encoding plant cell wall degrading enzymes (CWDE)

The *Fg* genome comprises a large number of genes encoding hydrolytic enzymes, transporters, secreted proteins and multiple gene clusters associated with secondary metabolite biosynthesis (Scherm et al., 2013; Sieber et al., 2014). These enzymes, non-enzymatic proteins and secondary metabolites together are generally considered fungal pathogenicity factors and their deployment during *Bd* root colonization indicates the importance of these pathway for sustaining the infection process. Indeed, over 30% of DEGs encode secondary metabolism enzymes (SMEs), secreted proteins, carbohydrate-active enzymes (CAZymes) and transporters (Fig. 1A). The most enriched categories for significantly up-regulated genes were associated with carbohydrate hydrolytic pathways (Suppl. data 3). Functional annotations showed that over half of the *in planta* activated CAZymes displayed modular structures related to CWDE acting on wall polymers such as cellulose, hemicellulose, lignin and pectin (Fig. 1B, Suppl. data 4), indicating the utilization of carbon from plant cell walls is a prominent capability of *Fg* during the colonization of *Bd* roots.

### Genes encoding secreted proteins and putative effectors

Fungal pathogens produce many small secreted proteins or effectors to help facilitate host colonization. However, relatively little is known about potential *Fg* effectors. By comparing our DEGs with the previously defined *Fg* secretome (Brown et al., 2012) and using the EffectorP 2.0 software (Sperschneider et al., 2018), we identified 250 putative secreted proteins along with 65 predicted fungal effector encoding genes (Fig S1C, Suppl. data 4 and 5). Many of these secreted protein genes, including 189 induced ones (Suppl. data 5), encode putative lipases and peptidases predicted to perform hydrolytic functions (Fig. 1C). Among the 65 predicted effector-encoding genes, 52 were significantly up-regulated during *Bd* root infection (Suppl. data 5). Some of these DE genes were reported to encode effectors actively secreted by *Fg* during *in vitro* growth. For example, *FG05_04074* encodes a protein of unknown function detected in two different secretomes (Lu and Edwards, 2016; Yang et al., 2012). Other putative effectors with annotated functional domains also found in previous studies included two glycoside hydrolases (FG05_11037 and FG05_06466), a putative acetylesterase (FG05_11280), a putative endonuclease (FG05_03365) and a cerato-platanin family protein (FG05_10212) for which roles of protection against host defense have been proposed (Lu and Edwards, 2016; Quarantin et al., 2016; Yang et al., 2012). Interestingly, the top induced effector candidates were mostly with predicted enzymatic functions (Suppl. data 5). Function of these differentially regulated putative effectors, such as FG05_04735 encoding a putative hypersensitive response-inducing elicitor and FG05_02255 a LysM domain containing protein (Suppl. data 5), could be predicted in comparison with those containing similar structural domains from different fungal pathogens and generally associated with host penetration, spore dispersal, triggering plant defense responses, inhibiting chitin-induced immunity or protecting against plant lysis (De Jonge et al., 2010; Khan et al., 2016; Lo Presti et al., 2015; Marshall et al., 2011; Mentlak et al., 2012). Although exact functions of these genes up-regulated during infection are largely unknown, it can be speculated that these secreted proteins and putative effectors could benefit the fungus during the colonization of host roots.

### Genes encoding secondary metabolism enzymes (SME)

*Fg* is known to produce many secondary metabolites (SMs) during infection (Ma et al., 2013). In line with this, a strong induction of expression could be observed for genes encoding key signature enzymes (Fig. S1A), including the longiborneol synthase CLM1 (FG05_10397), butenolide synthase (FG05_08079), TRI5 and the terpenoid synthase DTC1 (FG05_03066) during infection (Suppl. data 6), suggesting that the corresponding products culmorin, butenolide, trichothecene and carotenoid may be the major mycotoxins delivered by the fungus to facilitate root infection. Among them, butenolide and trichothecene pathways are known to be co-regulated *in vitro* and *in planta* (Sieber et al., 2014). In contrast, down-regulation or very low *in planta* expression of other key SME genes, such as *PKS12*, *NPS2*, *NPS1*, *PKS4* and *PKS10* (Suppl. data 6), indicates that certain types of mycotoxins such as aurofusarin, ferricrocin, malonichrome, zearalenone and fusarin C might not be highly produced by *Fg* during root colonization. Aurofusarin does not affect wheat head infection by *Fg* (Malz et al., 2005), whereas ferricrocin and malonichrome have been shown to be important for pathogenesis-related development of *Fg* (Oide et al., 2014). The tailoring enzyme genes are usually clustered and co-regulated with the corresponding signature enzyme genes in *Fg*. These genes encode cytochrome P450s, oxidoreductases, acyltransferases and methyltransferases mainly involved in the SM pathway responsible for biosynthesis and modification of SM products (Sieber et al., 2014). Therefore, up-regulation of a large portion of tailoring enzyme genes found in this study is consistent with the regulation pattern of the signature enzymes (Fig. S1B).

### Genes encoding fungal transporters

Transporter encoding genes mostly induced during *Bd* root colonization comprised a large group within the DE gene list (Fig. S1C, Suppl. data 7). Indeed, the regulation of transporter genes, particularly those associated with carbohydrate and nitrogen uptake as well as the ATP-binding cassette transporters (ABC transporters), is often linked to fungal nutrient assimilation, sensing, defense and pathogenicity status in pathogenic fungi (Abou Ammar et al., 2013; Coleman and Mylonakis, 2009; Divon and Fluhr, 2007; Gardiner et al., 2013; Schuler et al., 2015; Struck, 2015; Yin et al., 2018). Interestingly, the major facilitator superfamily (MFS) transporters associated with phosphate (Pi) transport and multidrug resistance (MDR) were mostly down-regulated during root infection compared to media alone controls (Fig. S1C, Suppl. data 7). During colonization of maize stalk, *Fg* overcomes Pi limitation by up-regulating high-affinity Pi transporter genes *FGSG_03172* and *FGSG_02426* (Zhang et al., 2016). The observation here that expressions of these genes during *Bd* root infection significantly reduced indicates a relatively rich root Pi environment under the experimental conditions and the time point examined. The down-regulation of MDR transporter genes suggested that they are possibly not essential for *Fg* to resist against root-derived anti-fungal compounds or self-derived toxins. The elevated expression of a large number of MFS-type carbohydrate transport genes together with the induction of PCWDE genes indicate that *Fg* preferentially utilizes carbon to accomplish the infection cycle and a state of glucose depletion may exist at the examined stage. The top induced ABC transporters (Suppl. data 7) exclusively belonging to the ABC-G type transporters are known to be associated with self-protection, possibly by effluxing of antifungal compounds in many pathogenic fungi (Coleman and Mylonakis, 2009). Thus, it is likely that these ABC transporters together with the MFS family multidrug resistance transporters could contribute to the virulence and fitness of *Fg* by detoxifying plant defense compounds.

We also observed that genes encoding amino acid-related transporters such as the amino acid/polyamine/organocation (APC) family and the amino acid/auxin permease (AAAP) family transporters, which are the major nitrogen transporters, were differentially expressed during root infection. In contrast, no inorganic nitrogen transporter gene showed altered expression (Suppl. data 7). This suggests that *Fg* root infection requires plant-derived organic nitrogen sources, and is consistent with the finding that polyamines as well as their amino acid precursors are potent DON inducers in *Fg* and play important roles during head infection (Gardiner et al., 2010, 2009). Interestingly, the highest induced transporter (*FG05_02278*, over 10-fold logFC) gene encodes a putative APC family protein transporter involved in choline uptake. Choline was identified as one of the major fungal growth stimulators in wheat anthers and implicated in promoting *Fg* virulence (Strange et al., 1972). Thus, it is possible that choline, in addition to amino acids and their derivatives, is another major factor contributing to *Fg* root colonization.

### A small set of ‘core’ genes is activated during infection of different hosts and tissues by *Fg*

To obtain additional insights into *Fg* pathogenicity, we compared the *Fg* genes found to be induced during root colonization in this study with those previously reported to be induced during the colonization of other hosts or tissues (Brown et al., 2017; Harris et al., 2016; Lysøe et al., 2011; Zhang et al., 2012, 2016). These previous studies have reported several subsets of *in-planta* expressed *Fg* genes at multiple infection time-points and different disease development stages. To make a broader comparison, *Fg* genes that were commonly induced during infection at any of the studied time-points were selected. These included 3591 *Fg* genes induced during the infection of wheat heads (Brown et al 2017), 5061 genes expressed during the infection of wheat and barley heads and maize ears (Harris et al 2016) as well as 344 and 3066 genes induced during the infection of wheat juvenile coleoptiles (Zhang et al., 2012) and maize stalks (Zhang et al., 2016), respectively (Suppl. data 8). Through these comparisons, a total of 38 *Fg* genes commonly induced across all gene lists were identified (Fig. 2A, Suppl. data 9). Some of these genes were also differentially expressed between a FHB resistant and a susceptible wheat genotype (Pan et al., 2018). Unexpectedly, no mycotoxin- or pathogenicity-related SME genes were present among these 38 genes (Fig. 2B). An ABC transporter gene, *FG05_04580* (*FgABC1*), and its flanking neighbor *FG05_04581*, which encodes a transcription factor highly inducible by the mycotoxin zearalenone (Lee et al., 2010), were found among these common genes. Deletion of *FgABC1* causes reduced virulence of *Fg* on tested wheat tissues (Abou Ammar et al., 2013; Gardiner et al., 2013). Interestingly, *FgABC1* and *FG05_04581* homologs in the closely related pathogen *F. culmorum* were both highly induced by the antifungal compound tebuconazole (Hellin et al., 2018). This indicates that FgABC1 and FG05_04581 could be involved in self-protection against various defensive chemicals consistent with the observation that FgABC1 contributes to protection against the fungicide benalaxyl (Gardiner et al., 2013). Furthermore, most of these common genes encode non-SM enzymes such as CAZymes, peptidases and putative effectors. Among them, *FG05_03624*, a gene encoding a secreted xylanase, was previously shown to promote necrosis during *Fg* head infection (Moscetti et al., 2015). Protein homologs of several of these genes were also shown to be virulence factors in other fungal pathogens. For instance, FG05_00028 is homologous to metallopeptidases (MEP1), which were shown to be apoplastic effectors in *F. oxysporum* and *M. oryzae* (Jashni et al., 2015; Yan and Talbot, 2016). In *F. oxysporum*, the metallopeptidase FoMEP1 and the serine protease FoSEP1 act synergistically to cleave host chitinases and prevent their degradation of fungal cell walls (Jashni et al., 2015). Indeed, a serine-type proteinase inhibitor encoded by *FG05_08012* found in our gene list, shows high similarity to FoSEP1 (Suppl. data 9). Another putative hypersensitive inducing elicitor FG05_04741 shows significant homology to the *Verticillium dahlia* effector PevD1, which was shown to be a secreted elicitor triggering host defense and cell death (Liang et al., 2018). We hypothesize that these putative *Fg* effectors may perform roles that are similar to those found in other fungal pathogens. Taken together, it can be hypothesized that a common set of *Fg* genes seems to play essential roles in *Fg* for successful colonization of different tissue types.

**Figure 2.**
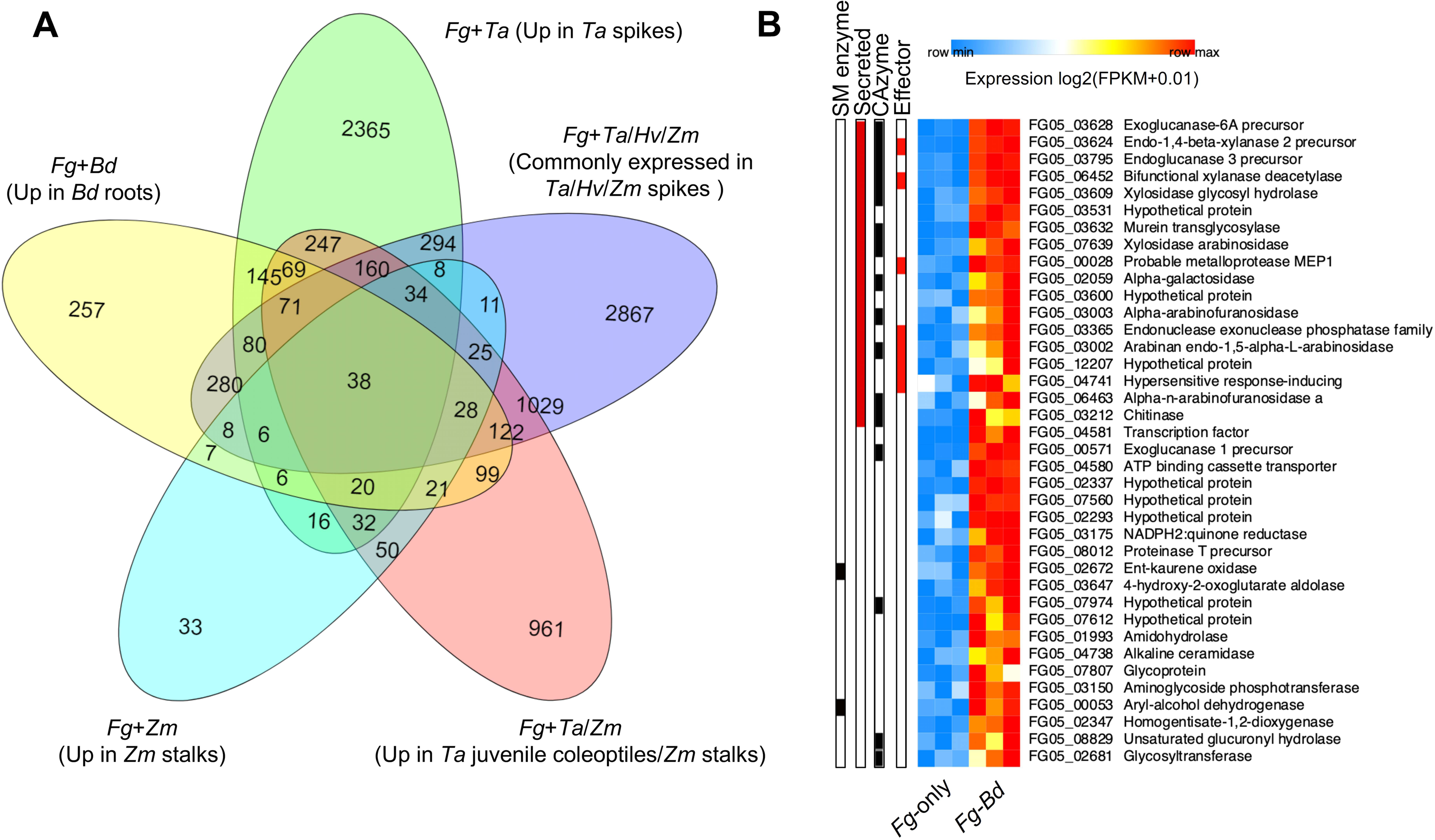
The number of *Fg* genes upregulated during *Bd* root infection were also induced or actively expressed during the infection of other tissues by *Fg*, suggesting that a core number of *Fg* genes could be broadly associated with infection (A) The Venn diagram showing how many *Fg* genes that were observed to be up-regulated during *Bd* root infection in this study were common to those upregulated or differentially expressed in different tissues of wheat (*Triticum aestivum*, *Ta*), barley (*Hordeum vulgare, Hv*) and maize (*Zea mays*, *Zm*). (B) The heatmap showing differential expression profiles of 38 core *Fg* genes identified from (A) between *in vitro* growth and during *Bd* root infection. FPKM values from three independent biological replicates were log2-transformed. Putative proteins encoded by these genes are shown on the right.

### *Fg* genes specifically upregulated in *Bd* roots

We have identified 257 *Fg* genes that were exclusively upregulated during *Bd* root infections (Fig. 2A). Functional category analysis showed a significant enrichment for genes involved in transmembrane transport and cellular import (FDR = 0.00334). Of 34 transporter encoding genes induced, 12 were predicted to be associated with carbon transport (Suppl. data 10). This supports the finding discussed above that carbon utilization by *Fg* plays a role during *Bd* root colonization. Three ABC-G and two MDR transporters (Suppl. data 10) found among the enriched transporters might be specifically associated with detoxification of *Bd* root defense compounds. Among the four predicted effectors induced in roots (Suppl. data 10), the putative host-necrosis inducer protein FG05_10212 was shown to be constitutively expressed during infections of wheat heads and *in vitro* and confirmed as an extracellular protein (Lu and Edwards, 2016). The induction of this effector might contribute to necrosis observed in the infected *Bd* roots. Notably, of the seven putative *Fg* PCWDEs whose transcripts were only induced in *Bd* roots, five use lignin as substrate (Suppl. data 10), suggesting that lignin-degradation by *Fg* in *Bd* roots. Overall, while some common infection strategies may be employed by *Fg* during infection of different hosts and tissue types, there appears to be also unique processes used based on the activation of specific *Fg* genes during root colonization.

### Partially shared infection strategies may be used by *Fg* and its sister species *Fp* during above- and below-ground infection of *Bd*

Previously, above-ground responses to the infection of *Bd* seedlings by *F. pseudograminearum* (*Fp*), another fungal species that is highly similar to *Fg* at the whole genome level (Gardiner et al., 2018) and was previously considered to be the same species as *Fg* (Kazan and Gardiner, 2018b), have been investigated (Powell et al., 2017). Both *Fg* and *Fp* show highly similar infection patterns on *Bd* (Fitzgerald et al., 2015). In addition, most genes are located in similar genomic regions in both fungi, whereas only a few species-specific genes, which could not be revealed by syntenic analysis, were found in genomic locations displaying high SNP densities (Gardiner et al., 2018). To further explore organ specificity of *Fg* infection on the same host, we compared the transcriptome of *Fg* with that of *Fp* (NCBI accession no. SRR3695327), during above ground infection of *Bd* at the same time point. Only *Fg* and *Fp* orthologous genes, which could be mapped to both of the genomes and share an identity of ≥99% were retained in this comparison. This stringent cut-off allows comparisons of only highly conserved genes that might be predicted to show similar pattern in expression and function in these two closely related fungal species.

In total, 1835 of the *Fg* DEGs were matched to *Fp* and formed a syntenic map (Fig. 3A and Suppl. data 11) consistent with the previously revealed genome structures (Gardiner et al., 2018). These genes were, in general, similarly expressed in *Fg* and *Fp* as indicated by the logarithmic transformed FPKM values (Fig. 3A). However, some variability in gene expression between the two species could be observed, particularly for genes found on chromosome 2 where the *Fg* orthologs tend to be preferentially expressed, as reflected by the expression heat map (Fig. 3A). Parts of chromosome 2 were previously identified as regions of the *Fg* genome that are rapidly evolving (Sperschneider et al., 2015). By manual curation, we selected and annotated the top 20 variant genes (Fig. 3B) that included three MFS-type transporter genes and an acetate permease homolog as well as several putative defense associated genes encoding a cell-wall glycoprotein (FG05_03352), peptidases (FG05_08075 and FG05_08141), and glucosidases (FG05_03387 and FG05_08265). A Zn_2_Cys_6_ transcription factor (FG05_03727), which shares the highest similarity to the yeast multidrug and oxidative stress resistance regulator STB5 (Larochelle et al., 2006), was identified. Accordingly, we found several oxidative stress responsive genes encoding a molybdopterin oxidoreductase (FG05_02880), NADH-flavin oxidoreductase (FG05_08077) and a putative flavohemoglobin (FG05_04458). Notably, *FG05_03914* encoding a putative isochorismatase (ISC) gene was only expressed in *Fg* during infection of *Bd* roots. ISC-like effectors in filamentous pathogens are conserved virulence factors that can subvert plant salicylic acid (SA) pathway and interfere with host immunity (Liu et al., 2014). Resistance to biotrophic and hemi-biotrophic pathogens is usually conferred by the host SA pathway (Pieterse et al., 2012), and therefore, ISCs might be employed by *Fg* to support its hemi-biotrophic lifestyle to attenuate SA produced by the roots. The absence of *Fp ISC* transcription is consistent with the observation that host SA levels were not elevated in *Bd* plants at the early stages of infection (Powell et al., 2017). Host plants may activate tissue-specific defense signaling in response to below-and above-ground attacks (Lyons et al., 2015). To manipulate such defense responses, a fungal pathogen must evolve to a high flexibility for successful infection progressed in different host tissues. Despite the use of two fungal species, the data provided here may suggest shared infection strategies, including the interference with host defense, employed by *Fg* to support its belowground colonization.

**Figure 3.**
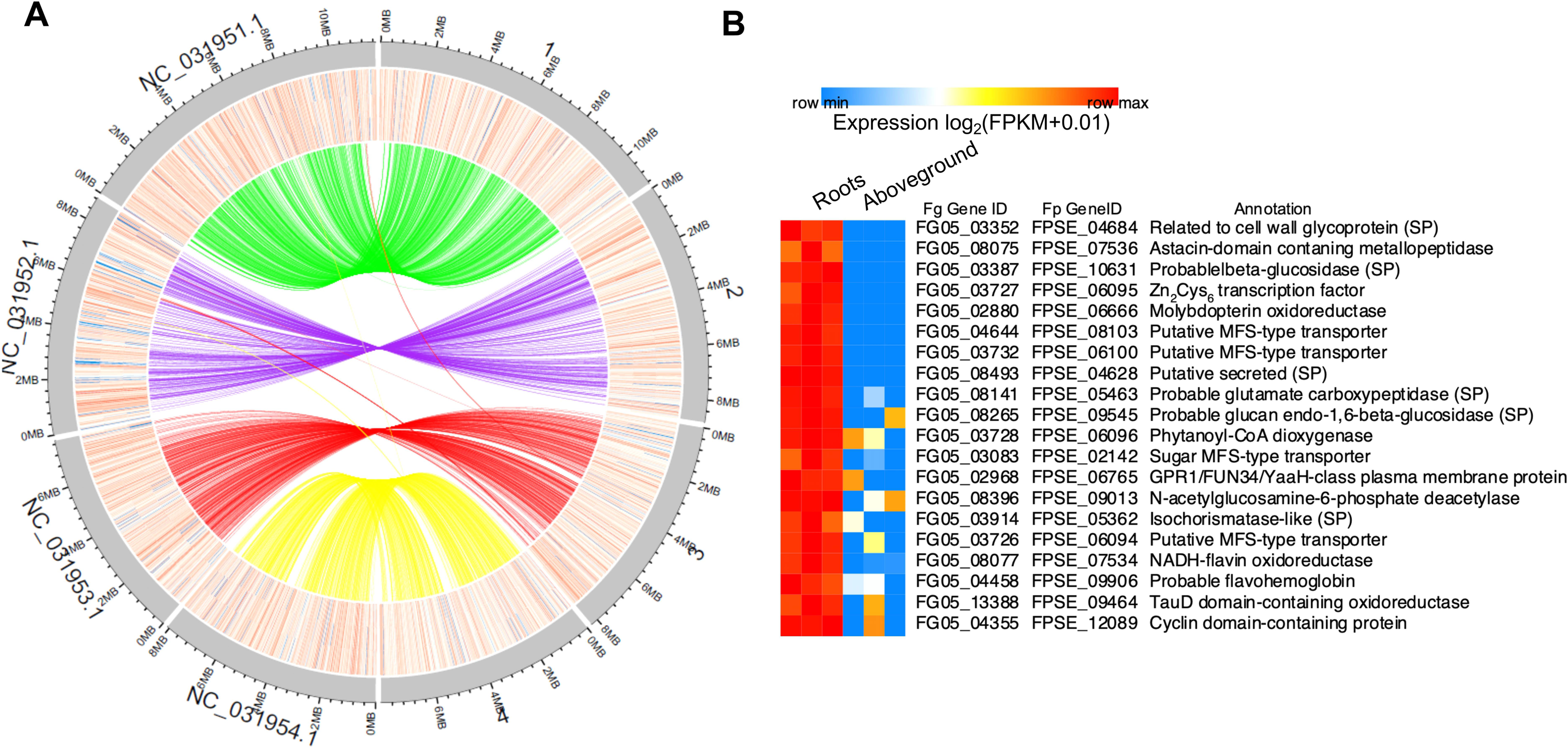
Genome-wide comparison of *Fg* and *F. pseudograminearum* (*Fp*) gene expression patterns during infection of *Bd* roots (this study) and aboveground tissues (Powell et al., 2017). (**A**) Chromosomal locations and syntenic relationship of highly conserved *Fg* and *Fp* genes. (1-4: *Fg* chromosomes; NC_031952.1-NC_031952.4: *Fp* chromosomes). (**B**) The top 20 genes with the most contrasted expression values were selected from chromosome 2 of *Fg* and *Fp* genomes, respectively. Heat maps in A and B display expression of genes as log2-transformed FPKM values from three biological replicates. Colour range is indicated as high (red) to low (blue).

### DON influences different fungal processes in *Fg* during *Bd* root infection

The trichothecene mycotoxin DON has been shown to significantly inhibit *Bd* root growth (Pasquet et al., 2016). However, to the best of our knowledge, there has not been any study examining the effect of DON on different fungal processes in *Fg*. Therefore, we conducted an RNA-seq analysis by infecting *Bd* roots with DON producing and deficient strains to determine *Fg* genes whose expressions are modulated by DON. We first focused on genes up-regulated in the DON deficient ΔTri5 mutant strain relative to WT. Of 973 genes differentially expressed between ΔTri5 and WT, 432 genes were expressed at higher levels in ΔTri5 (Fig. 4A, Suppl. data 12). These genes were subjected to functional enrichment analysis based on the MIPS FGDB (*Fusarium graminearum* Genome Database Functional Catalogue classification) (Güldener et al., 2006). This analysis showed that these genes are enriched for transport (FDR = 0.0016) of carbon-compounds, carbohydrates, and heavy metal ions, disease, virulence and defense (FDR = 0.005) and homeostasis of phosphate (FDR = 0.02). A relatively smaller portion of fungal pathogenicity and metabolism associated genes encoding CAZymes, SMEs, transporters and secreted proteins were induced in ΔTri5 relative to WT (Fig. 4B-C, Fig. S2 and suppl. data 12-13). Only metabolic pathway genes encoding methyl- and glycol-transferases as well as phosphate, lipid and polyamine transporters were preferentially induced (Fig. S2). Host-derived phosphates, lipids as well as polyamines may influence *Fg* infection in multiple host tissue types (Gardiner et al., 2010; Zhang et al., 2016). Their enhancement might be due to a positive feedback to the lack of DON to balance the fungal metabolism and cell structure in root proliferation. In addition, 28 CAZyme encoding genes were expressed higher in ΔTri5 than in WT and many of these were putative PCWDEs involved in lignin degradation (Fig. 4B). Lignin is one of the major barriers against fungal pathogens (Bhuiyan et al., 2009). Defense related or unrelated lignin content at fungal penetration sites might affect *Fg* intra-cellular progression (Zhang et al., 2016). In wheat roots, *Fg* colonization could be observed in lignin-rich vascular bundles (Bhandari et al., 2018; Wang et al., 2015). Therefore, the upregulation of lignin-degrading enzymes in ΔTri5 could be beneficial to the fungus to compensate DON deficiency and assist root colonization. In line with this, we observed reduced lignin deposition in roots colonized by ΔTri5 relative to the roots either mock-inoculated or colonized by WT *Fg* (Fig. 5).

**Figure 4.**
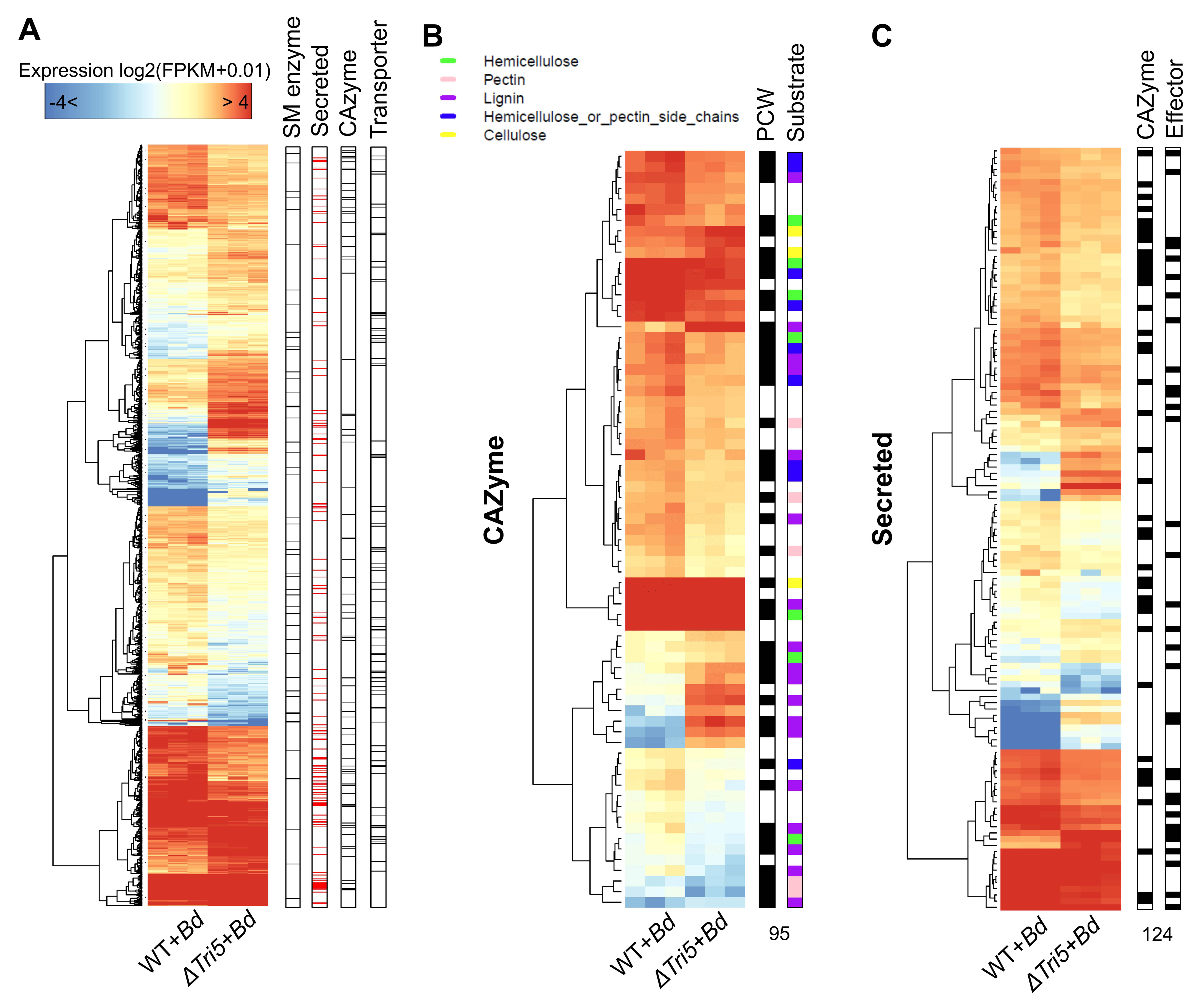
Global regulation of *Fg* genes in the *Tri5* mutant during *Bd* root colonization. (A) Expression of 974 *Fg* genes that were differentially expressed WT *Fg* and ΔTri5 during *Bd* root infection. Proteins encoded by DEGs were assigned to four functional categories as shown on the right. (B) Key features of 95 DEGs associated with carbohydrate active enzymes (CAZymes). Plant cell wall (PCW) degradation functions and substrate specificities were predicted as shown in the right-hand side. (C) Key features of 124 DEGs encoding putative secreted proteins. *Bd* roots inoculated with *Fg* WT and ΔTri5 strains were harvested at 5 dpi for the RNA-seq analysis. Transcript levels of *Fg* genes were log2-transformed and presented as normalized FPKM values. Heatmap colour range representing high to low expression levels is shown. Dendrograms show distance similarity of expression of each gene.

**Figure 5.**
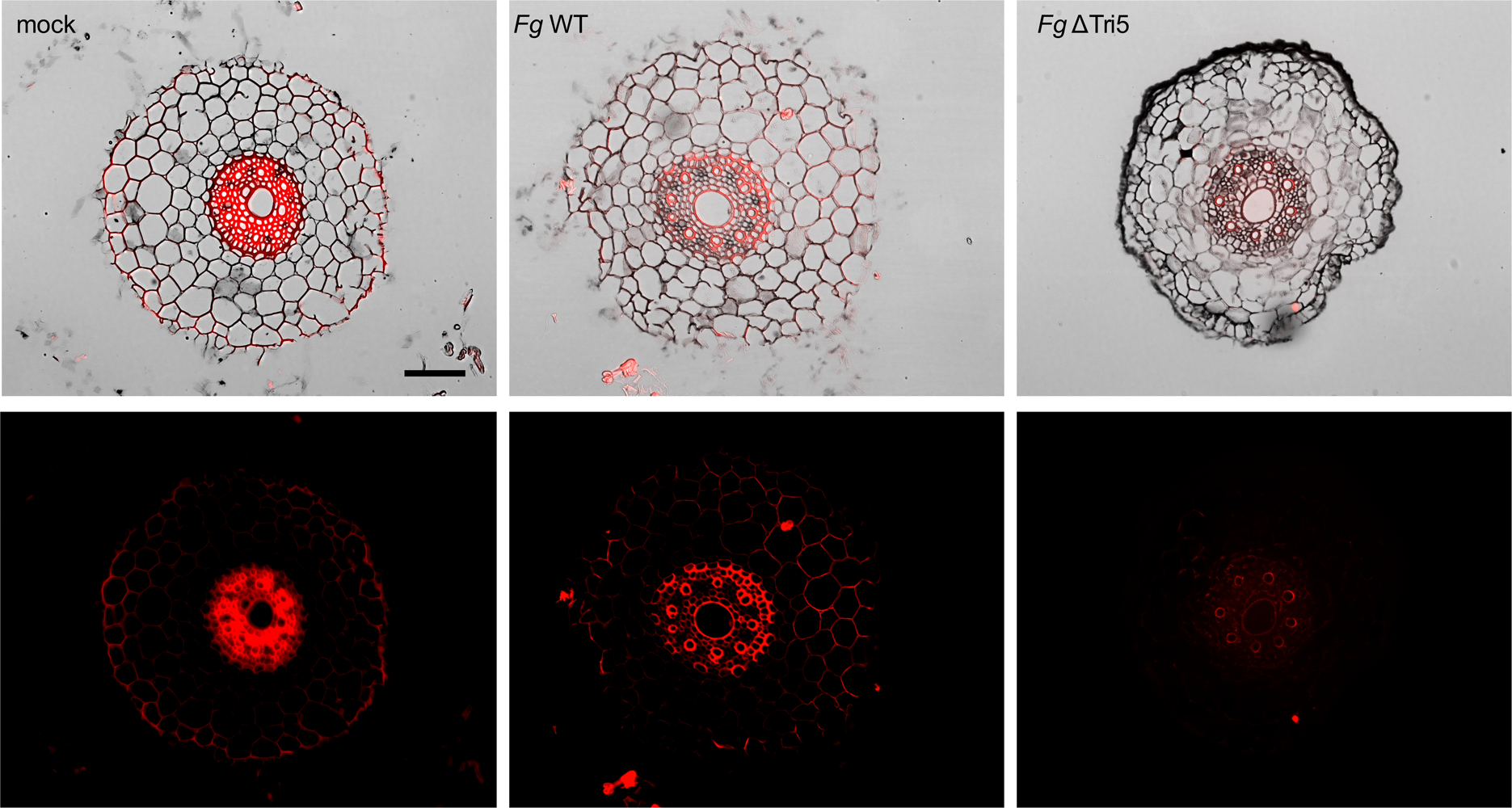
Lignin deposition in response to *Fg* infection is reduced in *Bd* roots inoculated with ΔTri5. *Bd* roots either mock-treated or inoculated with either WT *Fg* or ΔTri5 were harvested at 5 dpi. Red fluorescence signals (lower panels) indicate lignin deposition stained by Basic Fuchsin. Shown are representatives of at least 6 independent roots from three biological replicates. Images were taken using 561nm excitation and detected at 600–650 nm on a Zeiss Axio Imager M2 microscopy.

Seven of the 33 secreted protein genes induced in ΔTri5, including a pathogenesis-related protein 1 (PR1) homolog (FG05_03109) and a putative cutinase (FG05_03457), may be considered putative effectors. The most differentially regulated SME genes in ΔTri5 were tailoring enzyme genes encoding cytochrome P450s and oxidoreductases (Fig. S2A and S2B). Surprisingly, most DEGs with elevated transcripts levels in ΔTri5 during root colonization (405 out of 432 genes) were expressed either significantly lower in WT during *Bd* root infection than *Fg* only or remained unchanged comparing to WT *in vitro* (Suppl. data 12). Besides, we noticed that none of the above-mentioned core genes was reduced in ΔTri5 during infection, supporting the notion that these genes may contribute to infection more than others, independently of fungal DON production.

We next looked at the 541 significantly downregulated genes in ΔTri5 relative to WT in roots. The most enriched functional categories during root infection were C-compound and carbohydrate metabolism (80 genes, FDR = 0.00008), disease, virulence and defense (16 genes, FDR = 0.01), secondary metabolism (27 genes, FDR = 0.01), protein or peptide degradation (26 genes, FDR = 0.01) and transport facilities (42 genes, FDR = 0.03) (Suppl. data 14). In addition, we found two sets of adjacent genes *FG05_02297-FG05_02309* and *FG05_08077-FG05_08084* that showed reduced expression in ΔTri5. Of these two sets, the latter genes, which are part of the mycotoxin butanolide cluster, shared similar regulation pattern with the *Tri (tricothecene)* cluster genes during infection of wheat heads (Boedi et al., 2016). In the saprophytic fungus *Trichoderma arundinaceum*, the loss of trichothecene production likely contributed to an increase of fungal secondary metabolites (Lindo et al., 2019, 2018). Therefore, DON seems to affect fungal metabolism during *Fg* infection in *Bd* roots.

Previously, *Tri5* deletion was reported to lead to observable metabolic changes in *Fg* growing in rich medium, suggesting that DON might be linked to fungal physiology and development (Chen et al. 2011). In line with this, we also found that several differentially regulated TF genes in ΔTri5 during root infection were putative development-associated regulatory genes (Suppl. data 15). For example, FG05_08892 (MAT1-1-1) and FG05_05151 are known to be associated with sexual development (Kim et al., 2015), FG05_01139 (FgCBF1) is a predicted chromatin remodeling regulator (Guo et al., 2016), and FG05_03597 is homologous to *Aspergillus nidulans* FlbA, which is required for the control of mycelial proliferation and activation of asexual sporulation (Yu et al., 1996). However, when grown on MM, *Tri5* was barely expressed *in vitro*, and no DON could be detected by metabolic analysis (Fig. 6D). ΔTri5 also did not show any growth defects (Chen et al., 2011). Together, our results suggest that ΔTri5 may colonize the roots by utilizing a small set of genes not used by the WT fungus.

**Figure 6.**
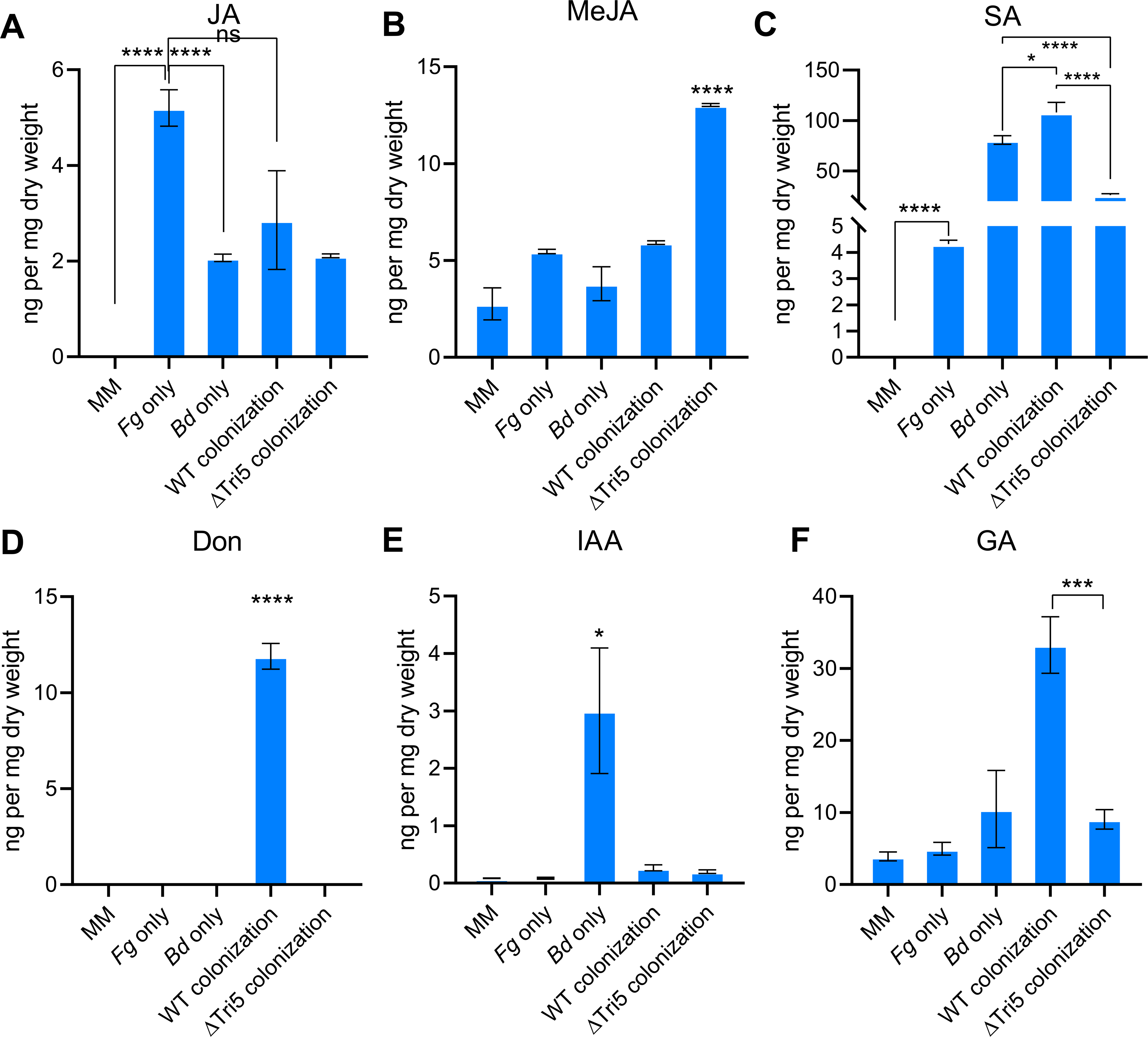
Quantification of selected phytohormones and DON in *Bd* roots inoculated with either WT *Fg* or the ΔTri5 mutant; jasmonic acid (JA) (**A**), methyl-jasmonate (MeJA) (**B**), salicylic acid (SA) (**C**), deoxynivalenol (DON) (**D**), indole-3-acetic acid (IAA) (**E**) and gibberellic acid (GA) (**F**) analysed by high-performance liquid chromatography. Independently grown fungal mycelium on minimal media (MM) (*Fg*-only), uninoculated *Bd* roots (*Bd*-only), as well as *Bd* roots inoculated with WT *Fg* or ΔTri5 infected roots were collected from MM at 5 dpi and subjected to metabolite extraction. Pure MM was used as background control. Metabolite quantifications were conducted according to the concentration-gradient derived standard curves of JA, MeJA, SA, DON, IAA and GA. Error bars indicate standard error of the mean based on six biological replicates, each comprising of 10 plants or 1 plate of fungal mycelium. Asterisks represent differences that were statistically significant (unpaired two-tailed t-test, *P< 0.05. ****P< 0.0001).

### DON is produced during *Bd* root colonization but does not act as a virulence factor

DON is a virulence factor during infection of wheat heads by *Fg* (Wang et al., 2020). However, it is unknown if this mycotoxin could also act as a virulence factor during infection of *Bd* roots. To determine this, the infection process was monitored using a *Fg* strain expressing Tri5-GFP fusion driving by the native *Tri5* gene promoter (Gardiner et al., 2009). Strong GFP signals could be visualized two days post-inoculation (dpi) in inoculated roots, indicating that the infection was progressing, and DON production was initiated (Fig. 7A and 7B). Consistent with this observation, *Tri5* was highly induced at 2 dpi and remained at high levels at later time points (3, 5, and 7 dpi) in the WT isolate (Fig. 7C). Previously, a temporarily similar infection pattern by *Fg* was also observed in wheat seedling roots (Wang et al., 2015). *Fg* root infection of wheat triggers induction of systemic defense responses in above-ground parts of the plant (Wang et al., 2018). In addition, DON preferentially inhibits root growth in wheat, *Bd* and Arabidopsis plants (Gatti et al., 2019; Masuda et al., 2007; Pasquet et al., 2016), and has been proposed to act as a major virulence factor in the early stages of *Fg* root infection (Wang et al. 2018). To assess the role of DON during root infection, a *Fg Tri5* mutant was used in inoculation experiments together with WT and Tri5-GFP strains. *Bd* roots infected by ΔTri5 exhibited levels of lesion development that were like those caused by WT and Tri5-GFP at 7 dpi (Fig. 7D). Therefore, while DON is a virulence factor in *Fg* during FHB of wheat and is highly induced in roots, *Bd* root infection by *Fg* seems to be independent of DON production. Indeed, various phytopathogenic phenotypes have been described for *Fg* DON deficient mutants (Boenisch and Schäfer, 2011; Cuzick et al., 2008; Jansen et al., 2005). For instance, altered levels of DON have been shown to inhibit plant apoptosis-like programmed cell death (PCD) induced by heat stress in Arabidopsis (Diamond et al., 2013).

**Figure 7.**
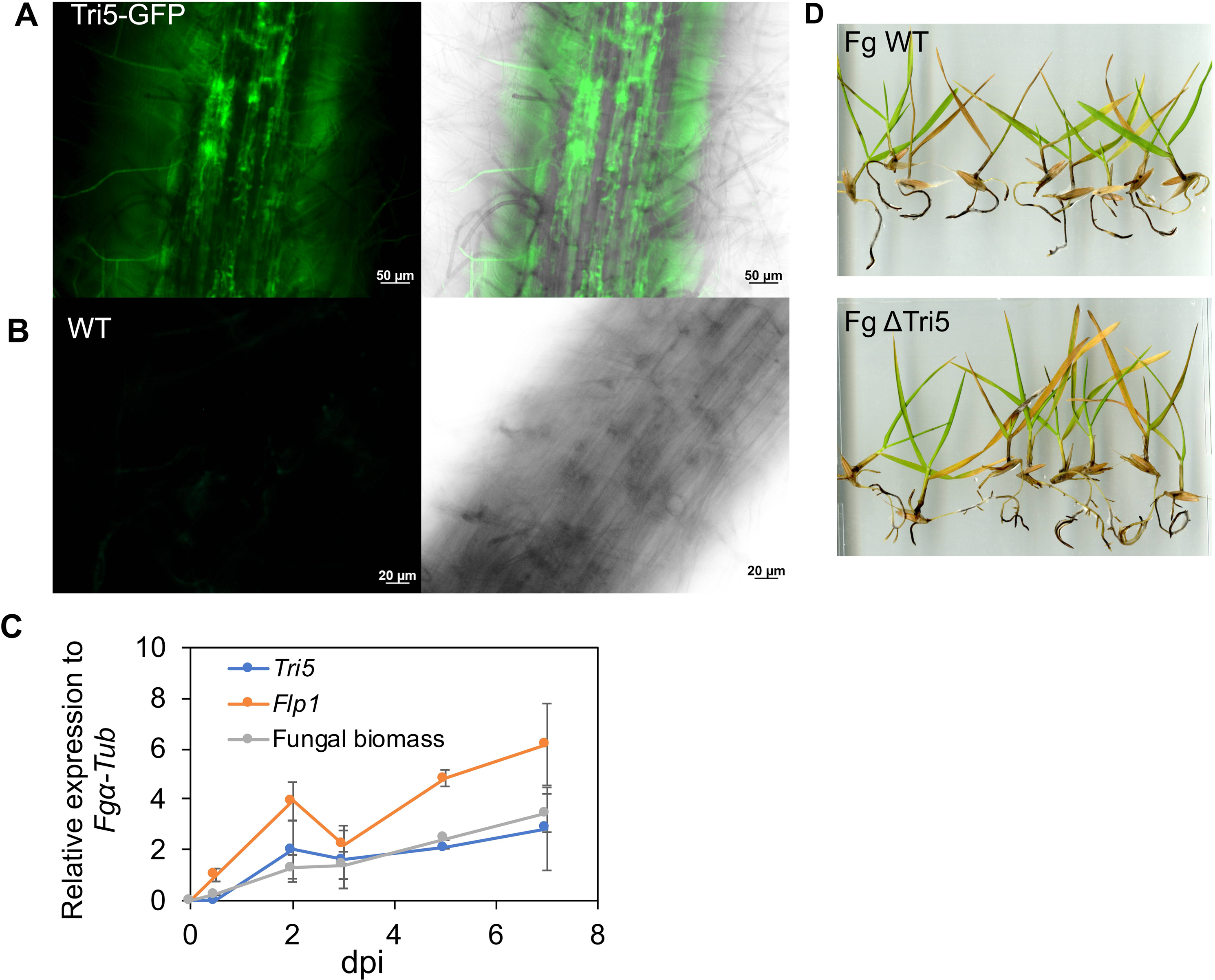
Deoxynivalenol (DON) produced by *F. graminearum* (*Fg*) during the infection of *B. distachyon* (*Bd*) roots does not contribute to lesion formation. Microscopic pictures were taken for *Bd* roots colonized by *Fg* strains after two days post inoculation (dpi). (**A**) DON is produced during the infection of *Bd* roots by *Fg*. Strong florescence signals could be observed in the mycelium of a *Fg* strain expressing a *Tri5-GFP* fusion construct driven by the native *Tri5* promoter. (**B**) No GFP signals could be detectable for WT *Fg* during root infection. **(C)** Transcriptional activation of *Tri5* confirms DON production in infected roots. Expressions of *Tri5*, *Flp1,* a lipase encoding gene previously identified as a virulence factor in *Fg* and a *Bd* ubiquitin-conjugating enzyme 18 gene (*UBC18*) used for fungal biomass measurements were quantified by RT-qPCR relative to the fungal *α-tubulin.* Root samples were harvested at 2, 3, 4, 5, and 7 dpi from three independent biological replicates each with at least 12 individual plants. (**D**) Deoxynivalenol (DON) does not contribute to lesion formation in *Bd* roots. Representative photos showing colonization of *Bd* roots with the WT *Fg* and *Tri5* deletion mutants.

### Phytohormone dynamics during *Fg* colonization of *Bd* roots

It is becoming increasingly evident that plant pathogens interfere with phytohormone pathways by producing plant hormones (Kazan and Lyons, 2014). However, pathogen-produced phytohormones have rarely been examined during root infections. We therefore next examined the transcriptome of *Fg* during *Bd* root infection, coupled with metabolic analyses, to determine putative phytohormone associated genes in *Fg* and their potential involvement in *Bd* root colonization.

### JA produced by *Fg* is not associated with *Bd* root colonization

The oxylipin hormone jasmonic acid and its derived metabolites collectively known as jasmonates (JAs) are derived from lipid peroxidation and can affect both host and fungal physiological processes (Tsitsigiannis and Keller, 2007). Fungal oxylipin biosynthesis is catalyzed by lipoxygenases (LOXs) (Fischer and Keller, 2016). In *F. oxysporum*, FoxLOX was found to exhibit a multifunctional activity in oxylipins production, thus proposed to possess a function in JA pathways (Brodhun et al., 2013). Interestingly, we noticed that the *Fg* homolog of FoxLOX, *FG05_05046*, was expressed during *in vitro* growth, and highly induced during root infection (4.8 fold, Table S1). JA-regulated defenses in plants can be interrupted by pathogen-derived hormone analogs (Caarls et al., 2017; Patkar and Naqvi, 2017). The *Fg* genome does not contain a homolog of the *M. oryzae antibiotic biosynthesis monooxygenase* (*Abm*), which converts host-derived JA into 12-hydroxyjasmonic acid (12OH-JA), thus attenuating rice blast disease resistance (Patkar and Naqvi, 2017). The Arabidopsis 2OG oxygenases (JOXs) are responsible for JA hydroxylation (Caarls et al., 2017; Smirnova et al., 2017). We identified ten homologous of Arabidopsis JOXs in *Fg* (Table S1). Of these, *FG05_08081* and *FG05_02301*, whose protein products share 22-26% identity to JOXs, were induced by 8.3 and 3.9 fold, respectively, during root infection. *FG05_08081* is present in the butanolide biosynthesis gene cluster, members of which were also significantly upregulated during root infection (Suppl. data 2). However, whether FG05_08081 and FG05_02301 are involved in JA degradation requires further analyses.

JA-associated host defense against *Fg* has been studied during FHB development. Inhibition of JA by DON at the bottom of wheat florets promotes fungal progression through rachis notes (Bönnighausen et al., 2019). Furthermore, late activation of JA signaling during FHB has been proposed to correlate with a necrotrophic transition of *Fg* (Ding et al., 2011). To determine if JA levels change during root infection, we quantified JA levels in *Bd* roots either mock-treated or inoculated with WT *Fg* or ΔTri5 strains by high performance liquid chromatography (HPLC). We found that 5 ng/mg dried material of JA was produced in *Fg* mycelia grown on MM (Fig. 6A). However, JA levels found in infected and control roots were much lower than 5 ng/mg and did not display any significant difference (Fig. 6A). Interestingly, however, higher levels of JA derivative methyl-JA (MeJA) were detectable in the roots infected by ΔTri5 than those infected by WT (Fig. 6B). Thus, DON seems to inhibit MeJA production in the infected *Bd* roots. This is consistent with the observation in wheat heads where MeJA levels in the ΔTri5-infected tissue were less than those infected with WT *Fg* (Bönnighausen et al., 2019).

### SA may synergistically interact with *Bd* root defense-related metabolic pathways

SA is a major defense hormone typically associated with plant defense against biotrophic pathogens (Glazebrook, 2005). JA and SA accumulate at different basal levels in various wheat cultivars and antagonistically fine-tune host defense responses (Powell et al., 2017). Therefore, we next focused on *Fg* responses to SA. Although the SA-pathway may be involved in host basal resistance against *Fg* (Ding et al., 2011; Makandar et al., 2010), SA-associated systemic acquired resistance (SAR) played no role in FHB resistance (Li and Yen, 2008). During infection of *Bd* seedlings by *Fp*, SA biosynthesis was induced (Powell et al., 2017). However, the function of SA during root infection by *Fg* remains elusive.

In Arabidopsis, SA biosynthesis during pathogen infection mainly relies on the intermediate chorismate processed by isochorismate synthase I (ICS1), the GH3 acyl adenylase-family enzyme PBS3, and the BAHD acyltransferase-family protein EPS1 (Torrens-Spence et al., 2019). The phenylalanine ammonia lyase (PAL) pathway also mediates SA synthesis through the conversion of benzoic acid or coumaric acid, but only contributes to a small portion of total SA production (Wildermuth et al., 2002). Similarly, some bacteria can directly convert isochorismate to SA by isochorismate pyruvate lyase (IPL) (Serino et al., 1995). While the host is believed to be the source of SA production in various plant-bacteria interactions, whether there is a fungal origin of SA remains unknown. In the *Fg* genome, we found two ICS1 homologs FG05_05195 and FG05_12934. Of these, *FG05_05195* was lowly expressed (FPKM < 0.5), but *FG05_12934* exhibited constitutive and high transcript levels during both *in vitro* growth and infection of *Bd* roots (Table S1). *Fg* also has an *EPS1* homolog, *FG05_00237*, which was significantly upregulated during *Bd* root infection (Table S1). *FG05_09331* encodes a protein sharing about 50% identity with PAL. *FG05_09331* expression decreased by 2.2-fold in infected roots relative to *Fg* grown *in vitro*. No PBS3 and IPL homologs could be identified in *Fg*. While expression patterns of *ICS1* and *EPS1* homologs may coincide with observed SA production by *Fg*, the absence of a *PBS3* homolog indicates other components or pathways could be involved in the SA biosynthesis. To determine if SA is produced in *Fg-Bd* interactions, we extracted metabolites from *Bd* roots with or without *Fg* inoculation and quantified SA levels by HPLC. In *Fg* mycelia, there was also about 0.01 µg SA per mg dry material (Fig. 6C). Most SA was found in the roots inoculated with WT *Fg*, followed by uninfected roots and the lowest levels in the ΔTri5 infected roots (Fig. 6C), indicating a potential role for DON in regulating SA levels during root infection by *Fg* although it is difficult to estimate the exact contribution of *Bd* or *Fg* to the SA levels measured.

SA has a direct effect on *Fg* growth, most likely associated with active degradation of SA by fungal hydroxylases. In recent studies (Hao et al., 2019; Qi et al., 2019; Rocheleau et al., 2019), two proteins FGSG_08116 and FGSG_03657 have been characterized with a function in SA degradation. While transcription of both FGSG_08116 and FGSG_03657 can be induced by external SA, fungal virulence in wheat heads was only influenced by deletion of the former. The function of FGSG_03657 for SA degradation was not disabled in deletion mutants. Interestingly, *FG05_03657* (a.k.a. FGSG_03657) was exclusively induced during *Bd* root infection (Suppl. data 10), suggesting that it may have a role in regulating SA levels in infected roots. To determine if *Fg* could possess additional putative SA hydroxylase genes (Hao et al., 2019), we searched the *Fg* genome and identified 28 homologs of the SA sensor and degradation protein Shy1 from *Ustilago maydis* with FgShyC displaying at least 20% identity to Shy1 (e-value < 10^-5^) (Fig. S3). This similarity is much higher than the values reported previously (Hao et al., 2019; Rabe et al., 2013), where the expansion of these putative proteins in *Fg* was supported by phylogenetic analysis (Fig. 8A). In our current transcriptome, *FG05_03657* and additional ten genes encoding putative SA hydroxylases were significantly upregulated in *Fg* during root infection (Fig. 8B). Therefore, SA biosynthesis by the pathogen as well as the host can contribute to the expression and regulation of these genes.

**Figure 8.**
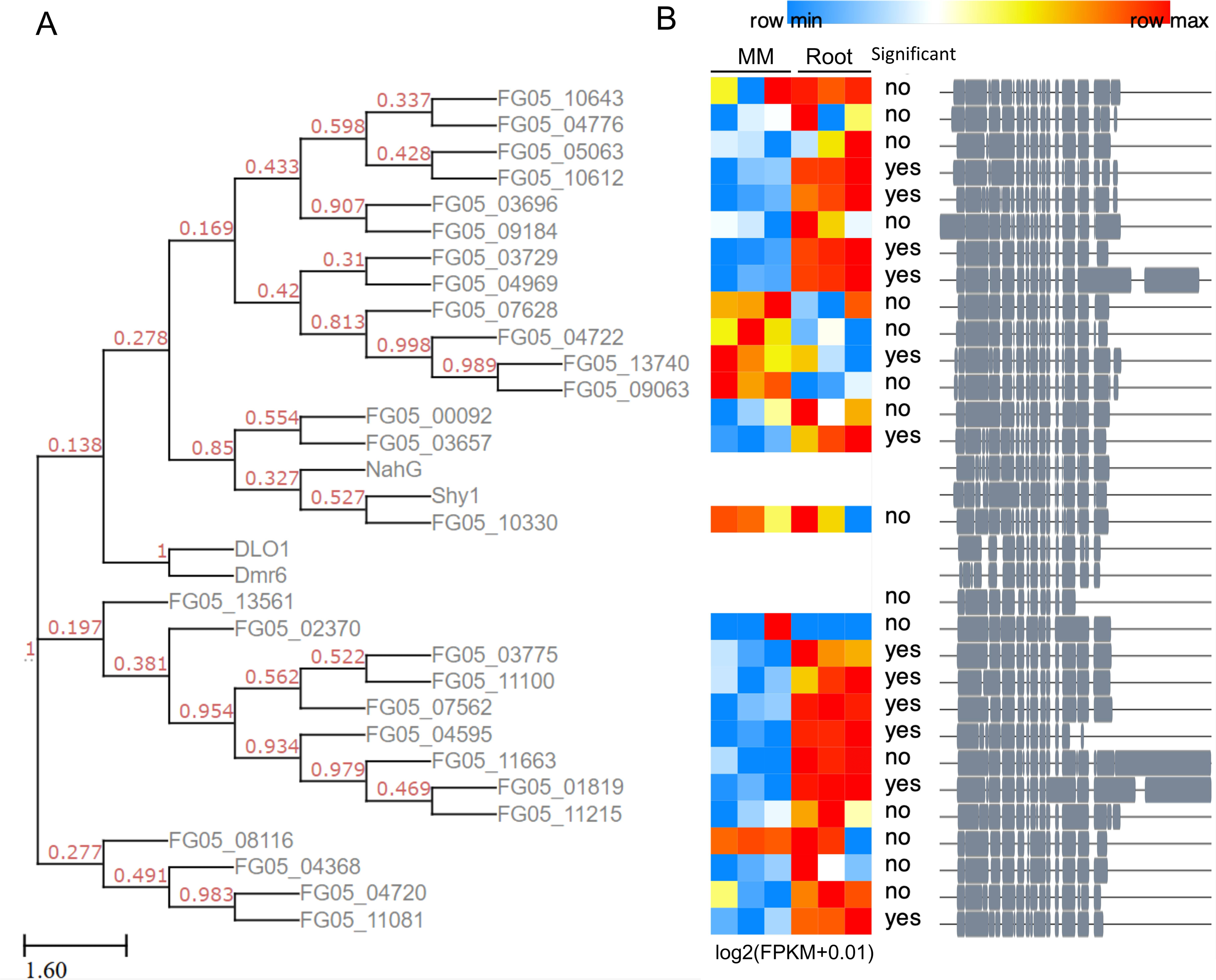
SA hydroxylase-like genes present in the *Fg* genome and their expression profiles during growth in culture or infection of *Bd* roots. (**A**) Phylogenetic unrooted tree of SA hydroxylase protein homologs of NahG, Arabidopsis Dlo1 (AT4G10500) and Dmr6 (AT5G24530) in *Fg*. The protein evolutionary models were tested using Neighbor-Joining inference and then referred to tree building based on Maximum Likelihood method. (**B**) Expressions of 28 SA hydroxylase candidate genes during *Bd* root infections or *in vitro* growth on minimal medium (MM). Heatmap shows expression levels of each gene displayed as log-transformed FPKM values of three replicates from low (blue) to high (red). *Fg* genes significantly induced during *Bd* root infection were marked as ‘yes’. An overview of concatenated alignment of all protein sequences was shown aside.

### Expression of genes involved in the biosynthesis of other phytohormones in *Fg*

### during *Bd* roots infection

In addition to JA and SA, other phytohormones such as gibberellins (GAs), auxins (IAAs), ethylene (ET), cytokinins (CKs) and abscisic acid (ABA) also participate in modulating host defense signaling (Pieterse et al., 2012). GAs can be synthesized by a number of *Fusarium* species but not by *Fg* due to the lack of a corresponding biosynthesis gene cluster (Cuomo et al., 2007). As a virulence factor, GA is restricted to the necrotrophic fungal pathogen *F. fujikuroi* (Wiemann et al., 2013) and is possibly involved in attenuating host JA signaling (Navarro et al., 2008). Under our inoculation conditions, GA was detected in the roots infected by WT *Fg* but not by the ΔTri5 mutant (Fig. 6F), indicating an endogenous GA production in *Bd* roots upon *Fg* infection and a potential positive effect of DON on root GA production. Similarly, GA accumulates in wheat heads infected by *Fg* (Bönnighausen et al., 2019), thus *Fg* seems to trigger a GA-dependent response in host roots that is similar to one observed in wheat florets (Buhrow et al., 2016). However, the association between DON and GA production requires further investigations.

Fungal genes encoding indole-3-acetaldehyde dehydrogenases (Iad) and tryptophan aminotransferases (IaaM) were thought to be responsible for IAA production (Reineke et al., 2008). A possible third pathway for auxin biosynthesis could be mediated by *Fg* genes homologous to YUCCA, a key enzyme involved in plant auxin biosynthesis (Mano and Nemoto, 2012). Interestingly, only one of the three *Fg Iad* gene homologs, *FG05_02773,* showed more than 4-fold induction in *Fg* during root infection as compared to *Fg* grown *in vitro* (Table S1). In contrast, two *IaaM* homologs and a *YUCCA* homolog were significantly downregulated in *Fg* inoculated roots (Table S1). Thus, the strong induction of *FG05_02773* might coincide with the production of the auxin indole-3-acetic acid (IAA) by *Fg*, which could thereafter compromise the host auxin pathway. Fungal auxin biosynthesis plays a role in pathogenicity of several pathogens (Chanclud and Morel, 2016). In *Fg*, auxin was proposed to be a virulence factor (Svoboda et al., 2019). Indeed, *Fg* is able to synthesize IAA but also sensitive to exogenous application of IAA and its biosynthetic intermediates (Qi et al., 2016). However, *Fg* infection strongly inhibited IAA levels in roots infected by *Fg* WT or ΔTri5 (Fig. 6E). This is contradictory to the findings in wheat where auxin levels increased during FHB (Wang et al., 2018). Factors such as host tissue types and hormone antagonists (Kazan and Manners, 2009) might be responsible for such differences.

*Fg* can exploit host ET signaling during colonization of both dicotyledonous and monocotyledonous plants and is believed to be capable of producing ET to counteract host defense pathways (Chen et al., 2009). However, rather than ET forming enzymes (EFE), *Fg* was thought to use pathways incorporating 1-aminocyclopropane carboxylic acids (ACC) as precursors for ET biosynthesis (Svoboda et al., 2019). Although an enzymatic function for two of the five ACC enzymes encoded by the *Fg* genome could be confirmed, fungal mutants for these genes showed no defect in pathogenicity on wheat (Svoboda et al., 2019). We looked at the expression of all these five genes, including the three annotated *ACC synthase* genes, *FG05_05184* (*ACS1*), *FG05_07606* (*ACS2*), *FG05_13587* (*ACS3*), and two *ACC deaminase* (*ACD*) genes, *FG05_02678* and *FG05_12669*, but found no differential expression during *Bd* root infections (Table S1), suggesting that pathogen produced ET may not be involved in root infection.

Biosynthesis of fungal cytokinins, which can be mediated by either one or both of the fungal transfer RNA-isopentenyl transferases (tRNA-IPT) and the Lonely Guy (LOG) enzyme, is associated with host immunity and nutrient modulation and maintenance of hemi-biotrophic lifestyles during infection (Spallek et al., 2018). Unlike many other *Fusarium* species, the *Fg* genome contains only one tRNA-IPT gene homolog (*FG05_09015*) (Sørensen et al., 2018). This gene was not differentially regulated (Table S1) and only moderately expressed (FPKM < 10) during both root infections and *in vitro* growth on MM.

Exogenous ABA has no effects on disease development, *Fg* toxin production or defense hormone levels in *Fg*-challenged wheat heads, but can promote fungal hydrolase and cytoskeletal reorganization genes induced early during infection and increase wheat’s susceptibility to FHB (Buhrow et al., 2016). The elucidation of fungal genes responsible for ABA production in the necrotrophic pathogen *Botrytis cinerea* and a few others has led to the hypothesis that a conserved ABA biosynthesis pathway exists in fungi (Lievens et al., 2017). In *B. cinerea*, such pathway involves four clustered genes *BcABA1*-*4* and a sesquiterpene cyclase gene *BcSTC5* (Izquierdo-Bueno et al., 2018). Fungal ABA was shown to act as a virulence factor in *M. oryzae*, which also harbors a direct ABA biosynthesis pathway but lacks a BsABA3 ortholog (Spence et al., 2015). Similar to *M. oryzae*, we could identify homologs of only BcABA1, 2 and 4 by BLASTp in *Fg*. In the current root transcriptome, none of these genes was differentially regulated (Table S1). Overall, while fungal GAs and IAAs might be associated with *Fg* root infections, it is unlikely that ETs, CKs and ABAs are involved in *Bd* root infection by *Fg*.

## Conclusions

The results presented here provide a detailed overview of root infection strategies employed by *Fg*, an important cereal fungal pathogen. The transcriptional regulation of pathogen metabolic pathways, virulence factors and signalling events during root infection show both unique and common features to those employed by *Fg* when infecting above-ground tissues. The mycotoxin DON, although not required for fungal virulence, produced during root infection appears to broadly affect various fungal processes and interplay with host responses. Expressions of several fungal stress and defence genes might help the pathogen to effectively deal with plant defence responses. In line with this, fungal JA, IAA and, in particular SA, seem to be used to interfere with root defenses. The findings presented in this paper will be useful for dissecting the mechanism of *Fg* belowground lifestyle and the development of novel plant protection strategies.

## Supporting information

Supplemental data sets

## Acknowledgements

We thank Di Xiao and Dr. Jonathan Powell for technical assistance. Yi Ding was the recipient of a post-doctoral fellowship from the Commonwealth Scientific and Industrial Research Organization Research Office.

## Conflict of interest

All authors declared no conflict of interest.

## Notes

### Competing Interest Statement

The authors have declared no competing interest.

