## Supplemental table and figures for "Transcriptome analysis reveals infection strategies employed by *Fusarium graminearum* as a root pathogen"

**Table S1. List of putative *Fg* phytohormone pathway associated genes.**

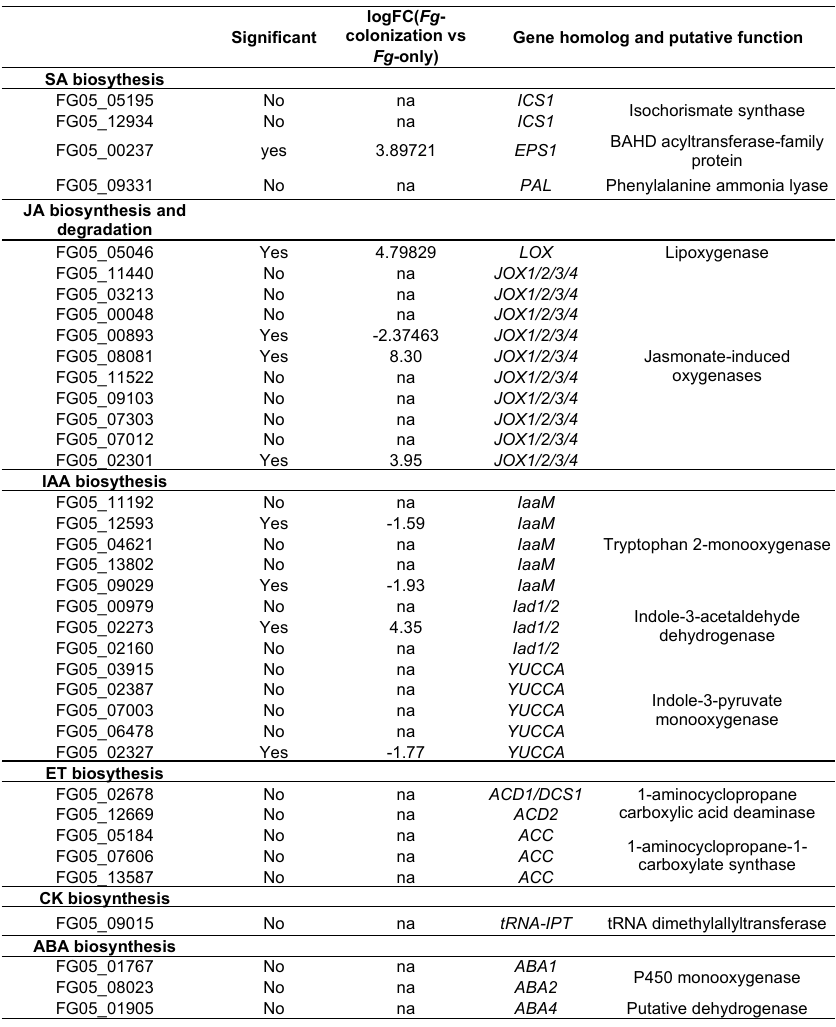

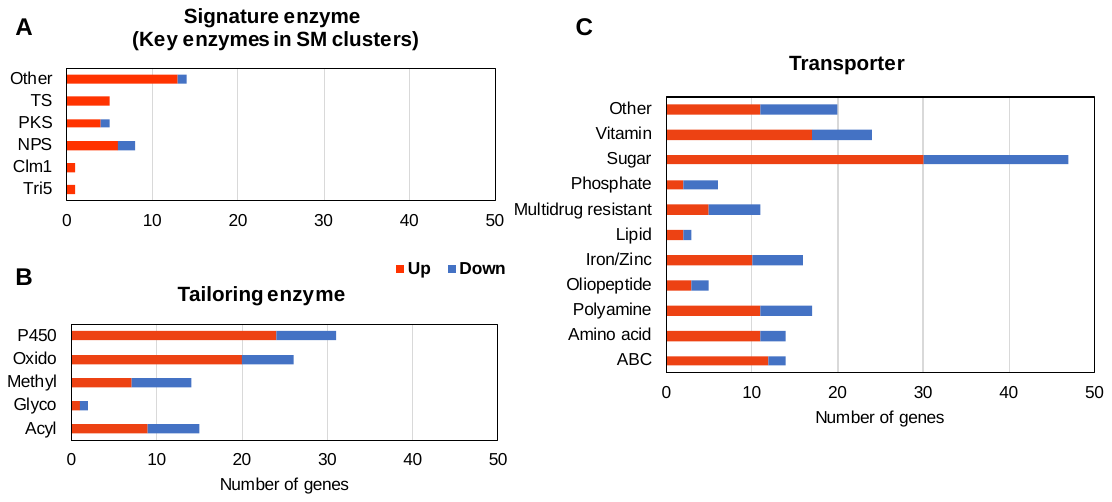

**Figure S1**. Number of *Fg* metabolic pathway genes differentially expressed between *in vitro* growth and in infected root samples. (A) DEGs encoding signature enzymes (key enzymes) involved in the secondary metabolite (SM) biosynthesis. TS: Terpenoid synthases, PKS: Polyketide synthases, NPS: Non-ribosomal peptide synthetases, Clm1: Culmorin synthase 1, Other: Putative enzymes. (B) DEGs encoding enzymes for modifications of metabolites. P450: cytochrome P450s, Oxido: oxidoreductases, Methyl: Methyltransferases, Glyco: Glycosyl-transferases, Acyl: Acyltransferases. (C) DEGs encoding transporters with predicted or unknown substrates.

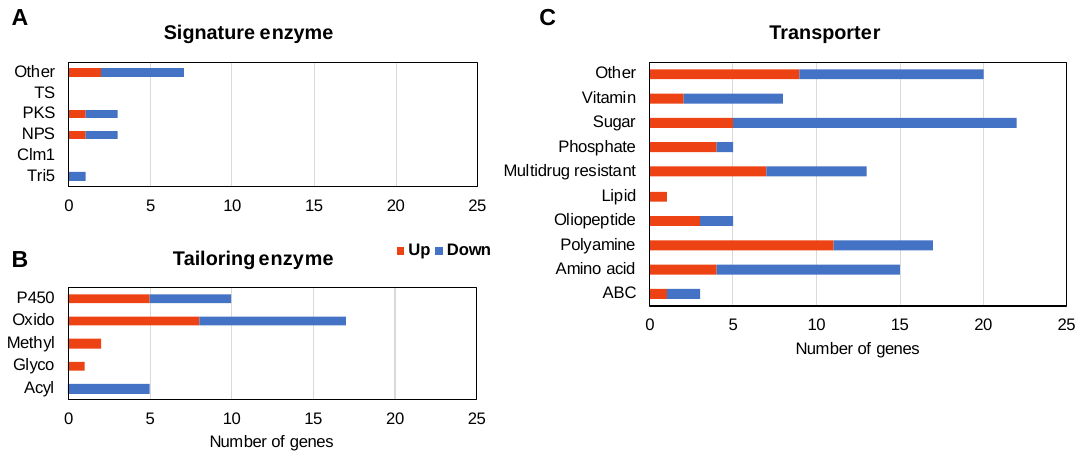

**Figure S2.** Differentially regulated fungal metabolic pathway genes in the *Tri5* mutant during *Bd* root colonization. (A) Number of up- and down-regulated *Fg* genes encoding signature enzymes involved in secondary metabolite biosynthesis (SM). (B) Number of up- and down-regulated *Fg* genes encoding enzymes putatively associated for modifications of metabolites. (C) Numbers of up- and down-regulated *Fg* transporter family genes with predicted or unknown substrates.

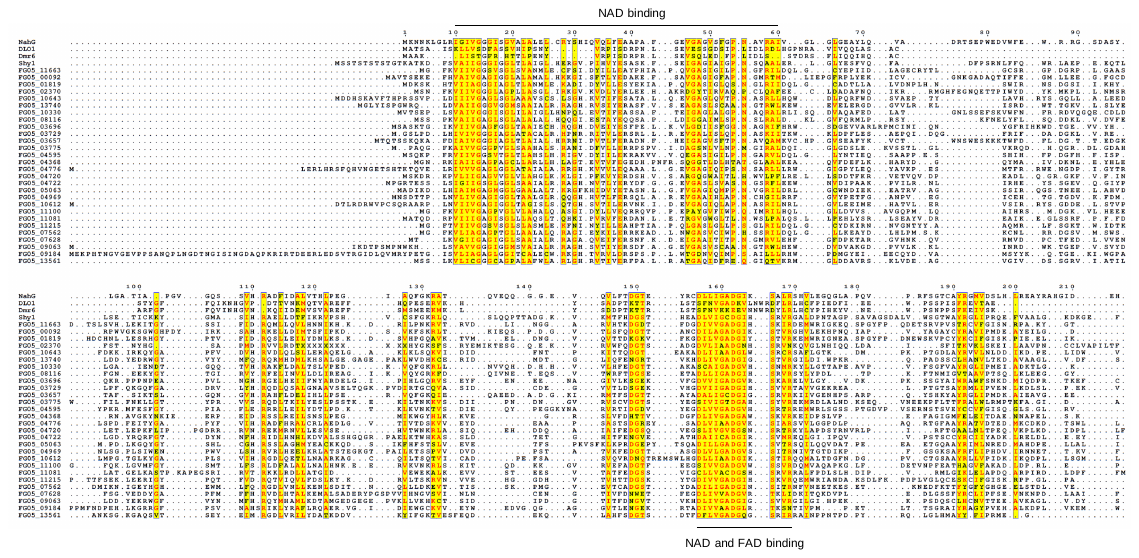

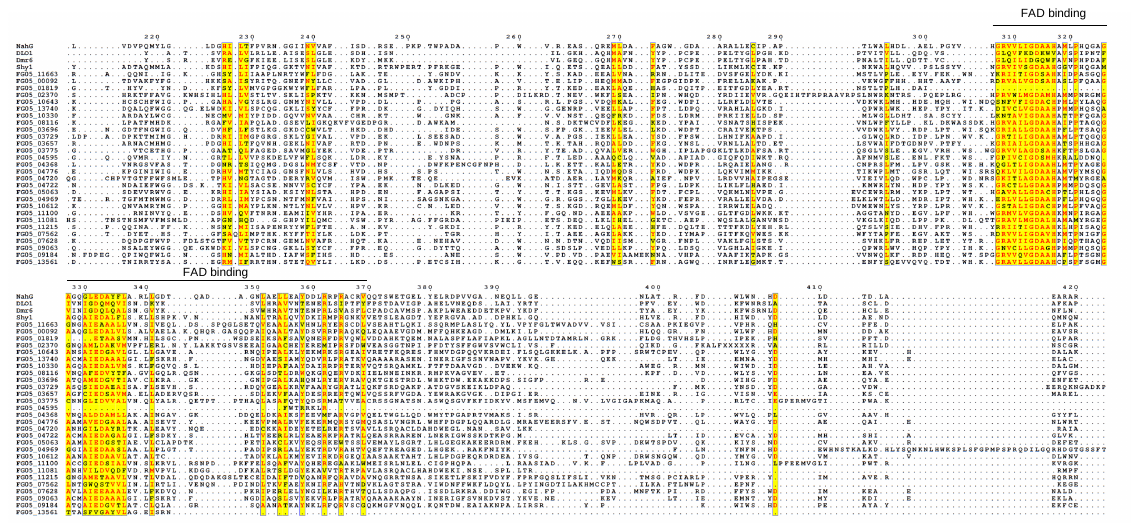

**Figure S3.** Sequence alignment of all SA hydroxylase gene candidates.

**Figure S4.** Normalization and statistics for the comparison of *Fg* and *Fp* transcriptomic datasets. For the analyses, raw FPKM values for all transcripts were log transformed.
